# Using mixed-effects modeling to estimate decay kinetics of response to SARS-CoV-2 infection

**DOI:** 10.1101/2021.02.22.432379

**Authors:** D. Bottino, G. Hather, L. Yuan, M. Stoddard, L. White, A. Chakravarty

## Abstract

The duration of natural immunity in response to SARS-CoV-2 is a matter of some debate in the literature at present. For example, in a recent publication characterizing SARS-CoV-2 immunity over time, the authors fit pooled longitudinal data, using fitted slopes to infer the duration of SARS-CoV-2 immunity. In fact, such approaches can lead to misleading conclusions as a result of statistical model-fitting artifacts. To exemplify this phenomenon, we reanalyzed one of the markers (pseudovirus neutralizing titer) in the publication, using mixed-effects modeling, a methodology better suited to longitudinal datasets like these. Our findings showed that the half-life was both longer and more variable than reported by the authors. The example selected by us here illustrates the utility of mixed-effects modeling in provide more accurate estimates of the duration and heterogeneity of half-lives of molecular and cellular biomarkers of SARS-CoV-2 immunity.

## Introduction

The calculation of decay rates is a common problem in the natural sciences. Combining direct experimental observation with mathematical modeling of decay kinetics allows for the calculation of half-lives that are many orders of magnitude larger than the observed time period. For example, the element Bismuth, usually thought of as stable, actually has a half-life of 20,000,000,000,000,000,000 years (7). This finding was made by observing the energetics and rate of alpha-particle decay (6), and mathematically modeling this data, which was gathered over an observation period that was (not surprisingly) far shorter than the published half-life.

A matter of slightly more pressing import at present is the calculation of half-lives of markers of immunity in response to SARS-CoV-2 infection. A number of reports in the popular press have put forward the idea that it is “too early to tell” how long the immune response to SARS-CoV-2 infection will last (8,9,10), as the observation period for most studies so far has been on the order of several months. However, the decay kinetics of humoral and cellular immunity are well understood processes, and tractable to mathematical modeling (11). Thus, careful model-based estimation of the duration of antibody, T-cell and B-cell responses can provide crucial insights into understanding the durability of immunological protection, which in turn will have a significant impact on the course of the pandemic in the coming years.

In this context, we can consider as an example the characterization of the immune response to SARS-CoV-2 infection by Dan et al. (*1*), published recently in Science. This study provides a comprehensive first look at the kinetics of memory T- and B- cell responses to SARS-CoV-2, and provides a contrast with the kinetics of the antibody response. The authors observe inter-individual heterogeneity in each immunological compartment examined, and comment that “…this heterogeneity means that long-term longitudinal studies will be required to precisely define antibody kinetics to SARS-CoV-2”.

To estimate half-lives, Dan et al., used a parsimonious approach, fitting either a linear or second-order polynomial to the log-transformed data pooled across patients, an approach often referred to as “naïve pooled fitting.” These pooled cross-sectional model fits provide a poor description of the data. The reported correlations correspond to average R^2^ values of 0.12, 0.18 and 0.06 for antibodies, memory B-cells and memory T-cells respectively (0.12 for the entire dataset), indicating that on average 12% of the total variability in each dataset is described by the model fit. Similarly, the *p*-values (while impressive) are misleading as they are only a test of the hypothesis that the independent variable changes in response to the dependent variable- all they tell us is that immunity changes over the observed time period.

While parsimony is always desirable in modeling, the approach used by the authors here leads to skewed estimates of both the population mean and the heterogeneity in the duration and half-life of immune protection. Notably, the naïve pooled approach cannot distinguish population heterogeneity from measurement error. Thus, the uncertainty in half-life estimation by this method cannot be used to draw inferences about population-level variability in protection against SARS-CoV-2 infection. In some cases, the authors did perform a longitudinal analysis, which they described as follows: “simple linear regression was performed, with t_1/2_ calculated from log2-transformed data for each pair”. However, linear “regression” to two data points is simply a line through those points (with zero residuals between data and model), and as such cannot distinguish between population heterogeneity and measurement error.

Mixed-effects modeling (MEM) is a more suitable methodology for dealing with longitudinal datasets such as these, where multiple correlated measurements are taken from each subject. Such an approach allows for more accurate and precise estimates of population heterogeneity and allows for it to be distinguished from the uncertainty in the population mean (*2*). This methodology is standard in drug development and clinical pharmacology for estimating the variability of pharmacokinetic and pharmacodynamic rate constants in longitudinal clinical datasets (*2*). Notably, MEM has been used to provide precise estimates of the long-term durability of antibody response after vaccination (*3*) or immunotherapy (*4*), as well as to describe the interrelation between different immunological markers (*5*).

## Methods

To isolate the effect of using mixed effects modeling versus naïve-pooled parameter estimation (rather than embarking on a model-building activity), we used the default Monolix (2020R1, Lixoft) settings and the same single-exponential model used by Dan et al (1) to fit PSV observations (*y_obs_*(*t*)):

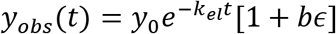

The parameter *y*_0_ is the estimated PSV titer at symptom onset, *k_el_* is the PSV titer exponential elimination rate (1/day), *t* is time in days post-symptom onset, and *b* is the estimated magnitude of proportional error between the model prediction and noisy observations. We performed nonlinear mixed effects modeling using Monolix Suite 2020R1 (Lixoft). The monolix-formatted dataset derived from the dataset provided in Supplementary Materials from (1), the monolix project and model files, and R code for generating the figures are included in the supplementary materials.

As we assumed inter-individual variability on both model parameters (*y*_0_ and *k_el_*), we obtained population estimates (Table 1) and individual patient parameter values (10 samples of each parameter drawn from the posterior parameter distribution for each patient). We then reported the histogram of all individual patient estimates for half-life 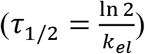 in Fig 1.

**Table 1:**
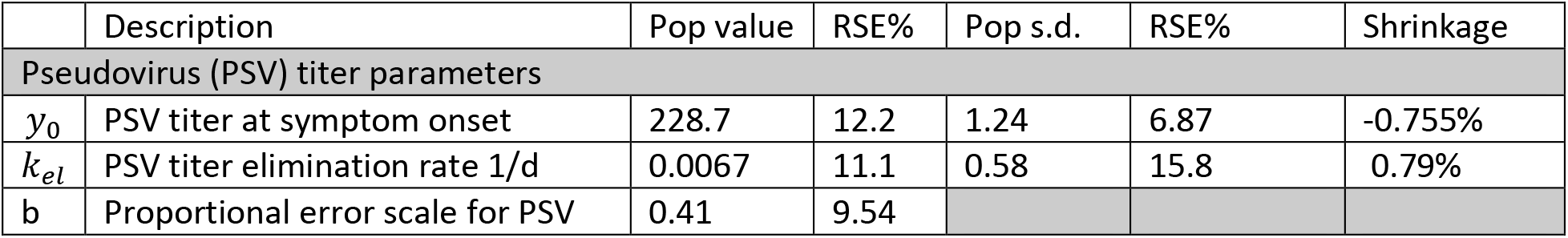
Population parameter estimates from MEM. RSE = Relative Standard Error. Population SDs are in lognormal space; a population SD of 1 corresponds to ~100% variability in the population for that parameter.

**Figure 1.**
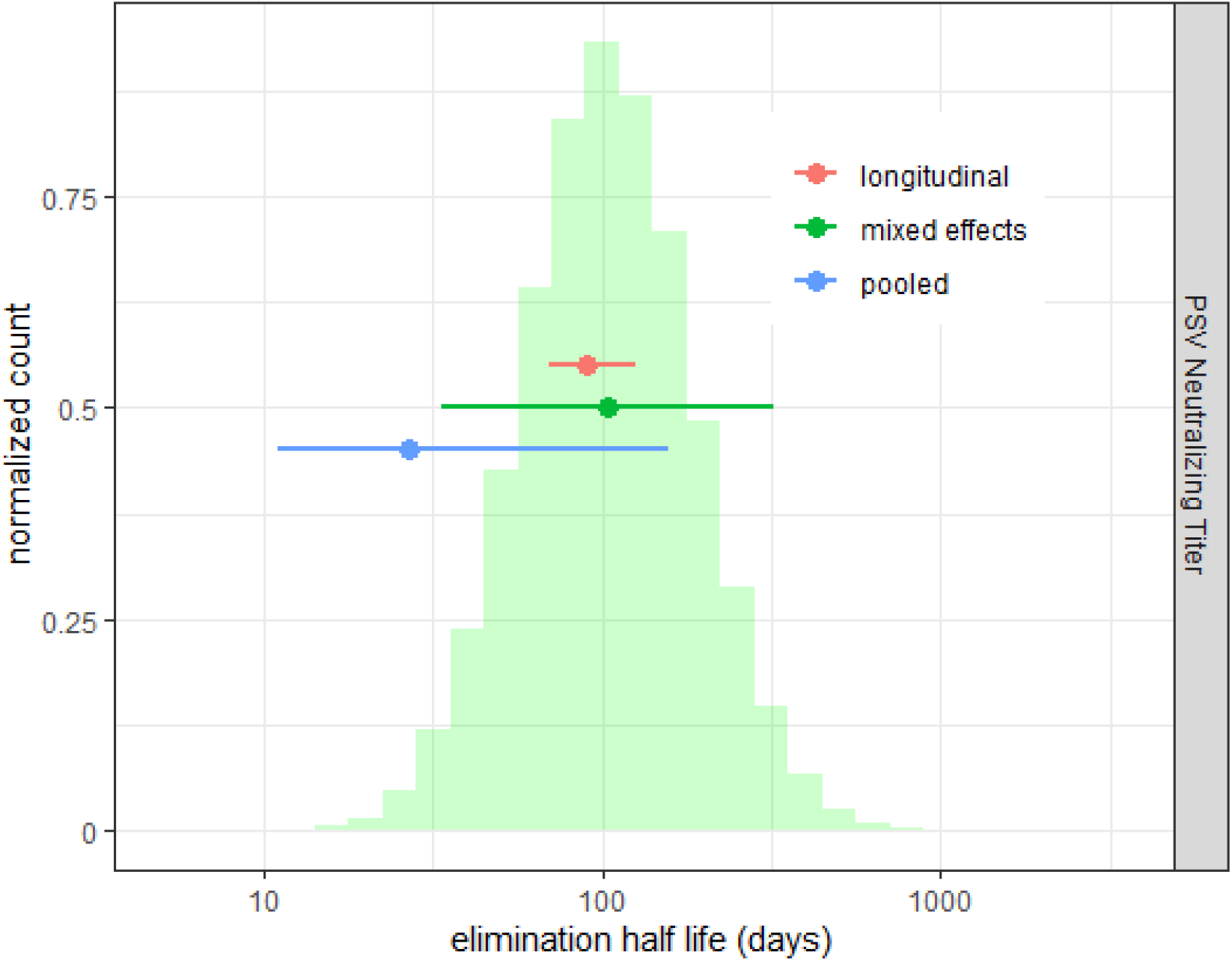
Distribution of half-life of pseudovirus neutralizing titer obtained from applying MEM for a single exponential model to the data from (1). Mean and 95% CI from pooled (blue), ‘longitudinal’ (red) analysis reported in (1) are shown as dot-and-whiskers plots. Green dot and whiskers indicate *population* median and 95% limits of estimated elimination half-lives from MEM.

To estimate the population distribution of time to loss of sensitivity, we simply calculated the time for each patient to decay to the limit of sensitivity *L* reported in Dan et al (L=20 for PSV): 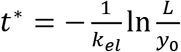. As a ‘posterior predictive check’ of this distribution of times to loss of sensitivity, we used the R (3.5.1) ‘survival’ package (2.42-3) in Rstudio (1.1.456, RStudio, Inc) to generate the empirical distribution of times as well as 90% bootstrapped confidence intervals. (Fig 2).

**Figure 2.**
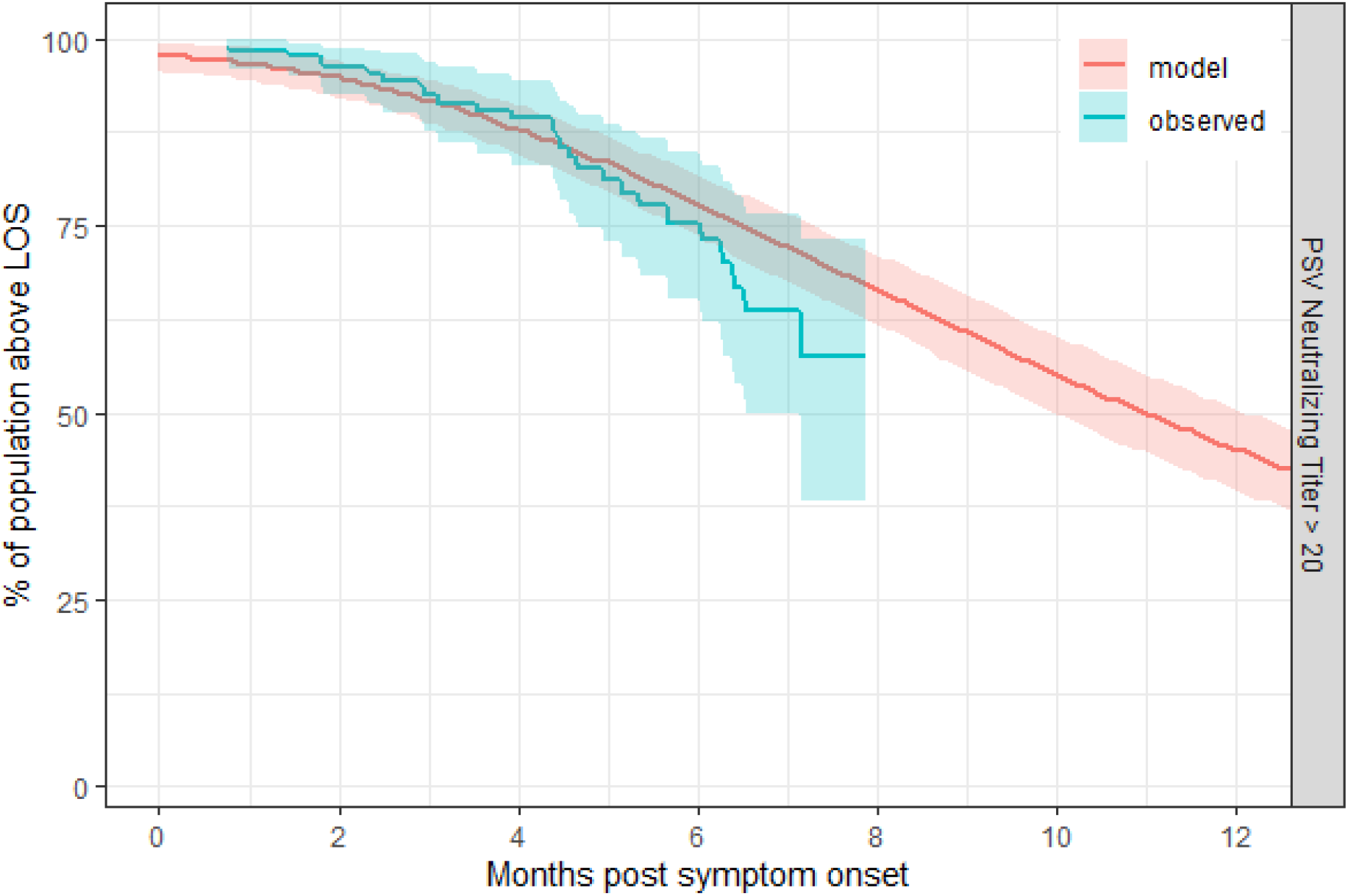
Estimated population distribution of time to return to LOS) for PSV. Blue time-to-event curve represents observed fraction above LOS (with bootstrapped 95% CI). Red curve represents estimated fraction using nonlinear MEM (with bootstrapped 95% CI).

## Results

In the illustrative example here, we have demonstrated the effect of using MEM rather than naïve pooled parameter estimation to fit a longitudinal dataset from the paper (pseudovirus neutralizing titer, or PSV). MEM estimates a median half-life of 100 days (95% population interval (95% PI): 33 −320d), compared to reported means of 27 days using naïve-pooled estimation (95% CI: 11-157d) and 90 days using longitudinal analysis (95% CI: 70-125d), as shown in Figure 1E in (*1*). Thus, the kinetics of PSV response is different than reported, as MEM reveals both a longer median half-life and a greater degree of heterogeneity. The latter difference is to be expected, as the 95% CI in the authors’ estimates reflects the uncertainty in the estimation of the mean half-life, whereas the 95% PI for MEM reflects the heterogeneity present within the patient population **(Fig. 1**).

We used the resulting fit from the MEM to estimate time taken for PSV to return to baseline (defined here as the limit of sensitivity, or LOS), finding that the median duration of return to LOS for pseudovirus neutralizing titer (PSV) was 332 days (95% CI: 39-1237 days), with 56% of the population predicted to return to LOS at 1 year (**Fig 2**).

## Discussion

The durability of the immune response to SARS-CoV-2 infection is a key determinant of the trajectory of the current pandemic. A number of reports have provided detailed longitudinal datasets describing this response, but have characterized their data using simplistic analyses that run the risk of misinterpretation.

As an example, in this report, we have selected one recent paper by Crotty et al (1), which has been the subject of substantial commentary in recent weeks. In the selected paper, the authors conclude that “immune memory in at least three immunological compartments was measurable in ~95% of subjects 5 to 8 months post-symptom onset, indicating that durable immunity against secondary COVID-19 disease is a possibility in most individuals”. The popular press has picked up on this particular scientific report, and it received extensive coverage, with numerous articles inferring from the author’s results that “the immune response to SARS-CoV-2 could last for years” (12,13,14). (We note that the conclusion in the popular press is stronger than the language used by the authors to describe their results, but the gist is the same in both cases).

For one marker (PSV) shown here as an example, our mixed-effects modeling analysis suggests a somewhat different interpretation. While some individuals can expect durable coverage against SARS-CoV-2 reinfection by this marker, PSV for approximately half of those infected by the virus will return to LOS by 12 months. MEM thus allows for a more precise estimate of the durability of the immune response, and also highlights the inter-individual variability, partitioning it away from experimental noise.

Using a modeling approach that is better suited to the datasets at hand has practical public-health implications. A growing number of cases of reinfection have been directly demonstrated using molecular phylogenetic analyses (15), and this functional data is consistent with population heterogeneity in the durability of the immune response. The existence of a subset of individuals who potentially lose immunological protection (and are vulnerable to reinfection) within 12 months is relevant both to the prospects for herd immunity and the public health strategy for this disease.

## Supporting information

R codes, analysis data set, and monolix files

## Author contributions

DB performed mixed effects modeling, co-wrote and edited the note. AC conceived the analysis, wrote the first draft and edited the note. GH and LFW provided input into the methodology and edited the note. MS and LY provided input into the methodology and conducted parallel but independent modeling of the data as a cross-check. All authors have read and approved the final manuscript.

## Data and materials availability

Models and code are available upon request from the corresponding authors.

---

**Figure S1:**
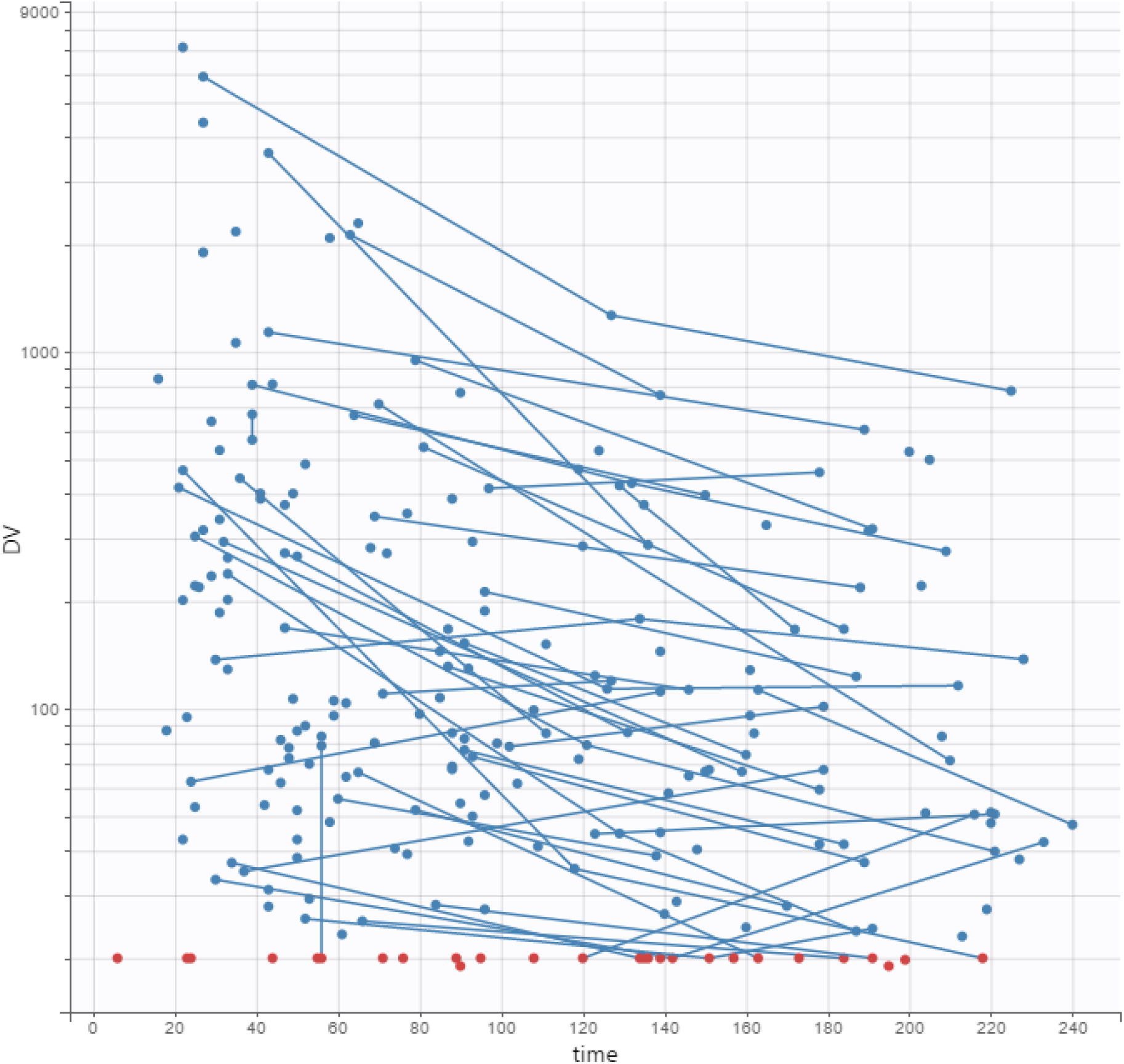
PSV titer data used in the analysis. Below limit of quantification data (BLOQ = 20) are shown as red dots. Data points from the same patient are joined with a blue line. Time is in days.

**Figure S2:**
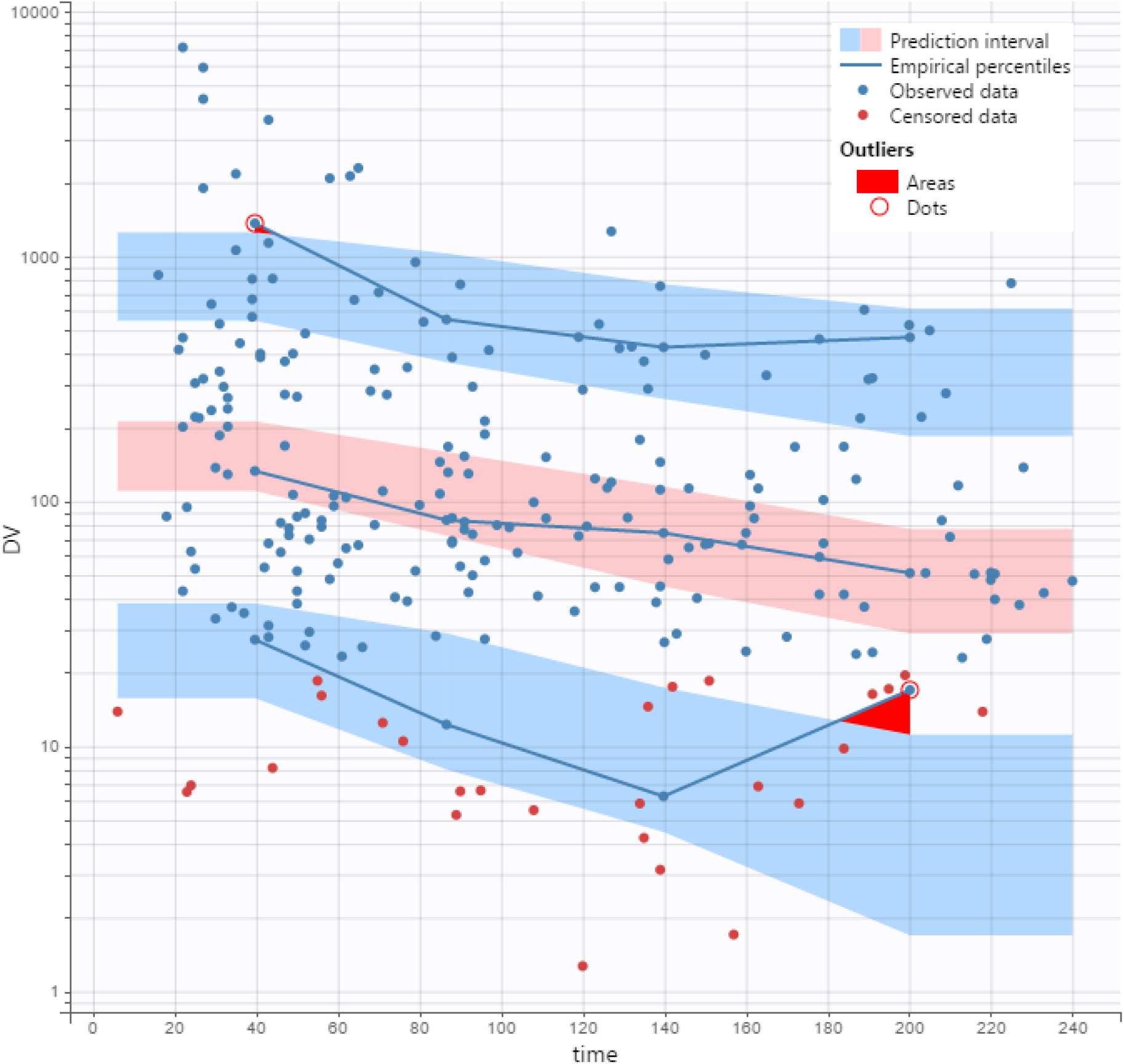
Visual Predictive Check (VPC) for Pseudovirus Neutralizing Titer (PSV). The blue and red bands represent the model-predicted 5, 50 and 95% population quantiles of the observations, while the blue lines represent the percentiles observed across the raw data. The red areas highlight deviation between the predicted and observed percentiles. The blue dots represent the observed data, and the red dots represent the simulated values of below limit of quantification (BLOQ) observations. The BLOQ values are simulated to avoid truncation artifacts in the VPC.

