## Supplementary material for "Using mixed-effects modeling to estimate decay kinetics of response to SARS-CoV-2 infection": R codes, analysis data set, and monolix files: postprocess_monolix.R

```
# plot distribution of stuff

rm(list=ls())

library(ggplot2)


dinm <- 365.25/12  # number of days in a month

runfun <- function(projname,Mnum=1,LOS,endpoint){

T <- read.csv(paste0('../monolix/',projname,'/IndividualParameters/simulatedIndividualParameters.txt'))

if (Mnum==2){T$TTel <- (T$Tin + log(T$kin*T$Tin/LOS)/T$kel)/dinm}
if (Mnum==1){T$TTel <- (-log(LOS/T$y0)/T$kel)/dinm}

T$Thalf = (log(2)/T$kel)/dinm

T$endpoint <- endpoint
T$Nsamples <- nrow(T)

return(T)

}

PSV <- runfun(projname='run015_PSV-neut-titer_M01-base',Mnum=1,LOS=20,endpoint='PSV Neutralizing Titer')
#Tcells <- runfun(projname='run013_CD4-spike-T_M01-base',Mnum=1,LOS=0.03,endpoint='Spike-specific CD4+ Tcells (%)')

#Tbig <- rbind(Tcells,PSV)
Tbig <- PSV

months <- c(6,12,18,24)
cusfun <- function(months){length(which(PSV$TTel < months))/nrow(PSV)}
halfun <- function(months){length(which(PSV$Thalf < months))/nrow(PSV)}
cprobs <- mapply(cusfun,months)
halfprobs <- mapply(halfun,months)
QPSV <- data.frame(months,cprobs,halfprobs)
QPSV$endpoint <- 'PSV Neutralizing Titer'

# cusfun <- function(months){length(which(Tcells$TTel < months))/nrow(Tcells)}
# halfun <- function(months){length(which(Tcells$Thalf < months))/nrow(Tcells)}
# cprobs <- mapply(cusfun,months)
# halfprobs <- mapply(halfun,months)
# QTcells <- data.frame(months,cprobs,halfprobs)
# QTcells$endpoint <- 'Spike-specific CD4+ Tcells (%)'

# QQ <- rbind(QPSV,QTcells)

QQ <- QPSV

ggplot(data=Tbig) + stat_ecdf(aes(x=TTel,color=endpoint),size=2) + 
  coord_cartesian(xlim=c(0,30))+ scale_x_continuous(breaks=c(6,12,18,24)) +
  scale_color_discrete() +
  scale_y_reverse(labels=function(x){100-100*x}) +
  xlab('Months post symptom onset') + 
  ylab(paste0('% protected')) + 
  geom_label(data=QQ,aes(x=months,y=cprobs,label=paste0(100-100*signif(cprobs,2),'%'),color=endpoint)) +
  theme(legend.position = c(0.8,0.8))

#quants <- c(0.025,0.25, 0.5, 0.75,0.975)
quants <- c(0.025, 0.5, 0.975)
sizes <- c(0.1,0.15,0.1)
hvals <- quantile(PSV$Thalf,quants)
QVPSV <- data.frame(quants,hvals,sizes)
QVPSV$endpoint <- 'PSV Neutralizing Titer'

# hvals <- quantile(Tcells$Thalf,quants)
# QVTcells <- data.frame(quants,hvals,sizes)
# QVTcells$endpoint <- 'Spike-specific CD4+ Tcells (%)'
# 
# QV <- rbind(QVPSV,QVTcells)

QV <- QVPSV

p.lb <- c(70,11,QV$hvals[1]*dinm)
p.mean <- c(90,27,QV$hvals[2]*dinm)
p.ub <- c(125,157,QV$hvals[3]*dinm)
p.y <- c(0.55,0.45,0.50)
analysis <- c('longitudinal','pooled','mixed effects')
endpoint<- c('PSV Neutralizing Titer','PSV Neutralizing Titer','PSV Neutralizing Titer')
pub.Thalf <- data.frame(p.lb,p.mean,p.ub,p.y,endpoint,analysis)
ymax <- 30000 

boo <- ggplot(data=Tbig) + stat_bin(aes(x=log10(Thalf*dinm)),binwidth=0.1,fill='green',alpha=0.2) + 
  coord_cartesian(xlim=log10(c(5,3500))) +  scale_x_continuous(breaks=c(1,2,3),labels=function(x){10^x}) +
  scale_color_discrete() + # scale_x_log10() + 
  xlab('elimination half life (days)') + 
  scale_y_continuous(breaks=ymax*seq(0,1,by=0.25),labels=function(y){signif(1*y/ymax,2)}) +
  ylab(paste0('normalized count')) + 
  #geom_vline(data=QV,aes(xintercept=log10(hvals*dinm)),color='blue') +
  #geom_label(data=QV,aes(x=log10(hvals*dinm),y=0,label=paste0(signif(dinm*hvals,2),'d')),color='blue') +
  #geom_label(data=QV,aes(x=log10(hvals*dinm),y=ymax,label=paste0(quants*100,'%')),color='blue') +
  theme_bw() + theme(legend.position = c(0.75,0.75),legend.title=element_blank()) + # theme(legend.position='none') +
  geom_point(data=pub.Thalf,aes(x=log10(p.mean),y=p.y*ymax,color=analysis),size=3) + 
  geom_segment(data=pub.Thalf,aes(x=log10(p.lb),xend=log10(p.ub),y=ymax*p.y,yend=ymax*p.y,color=analysis),size=1) + 
  facet_grid(rows=vars(endpoint))

print(boo)
  
```
