## Supplementary material for "Using mixed-effects modeling to estimate decay kinetics of response to SARS-CoV-2 infection": R codes, analysis data set, and monolix files: postprocess_monolix_bootstrap+PPC.R

```
# plot distribution of stuff with CI's

rm(list=ls())

library(ggplot2)
library(matrixStats)
library(survival)
library(data.table)
#library(survminer)

dinm <- 365.25/12  # number of days in a month

bootstrp <- function(projname,Mnum=1,LOS,endpoint,CIwidth = 0.95,Nboot=5000,tsim=seq(0,36,by=.1)){

#projname='run015_PSV-neut-titer_M01-base';Mnum=1;LOS=20;endpoint='PSV Neutralizing Titer'

T <- read.csv(paste0('../monolix/',projname,'/IndividualParameters/simulatedIndividualParameters.txt'))

if (Mnum==2){T$TTel <- (T$Tin + log(T$kin*T$Tin/LOS)/T$kel)/dinm}
if (Mnum==1){T$TTel <- (-log(LOS/T$y0)/T$kel)/dinm}

T$Thalf = (log(2)/T$kel)/dinm

T$endpoint <- endpoint


Nrep = max(T$rep)
uPAT = unique(T$id)
Npat = length(uPAT)

bootfun = function(Nrep,Npat){
booti <- sample.int(Nrep,size=Npat,replace=TRUE)  # which replicate to take for each patient
bootbase <- seq(0,Nrep*Npat-1,by=Nrep)
sampi <- bootbase + booti
ysamp <- T$TTel[sampi]
ec <- ecdf(ysamp)
return(ec(tsim))
}

foop <- replicate(Nboot,bootfun(Nrep,Npat))

QQ <- rowQuantiles(foop,probs=c((1-CIwidth)/2,0.5,1-(1-CIwidth)/2))
lower <- QQ[,1]
median <- QQ[,2]
upper <- QQ[,3]

Tout <- data.frame(tsim,lower,median,upper)
Tout$endpoint <- paste(endpoint,'>',LOS)
Tout$type <- 'model'

return(Tout)

}

boot.PSV <- bootstrp(projname='run015_PSV-neut-titer_M01-base',Mnum=1,LOS=20,endpoint='PSV Neutralizing Titer')
boot.Tcells <- bootstrp(projname='run013_CD4-spike-T_M01-base',Mnum=1,LOS=0.03,endpoint='Spike-specific CD4+ Tcells (%)')

Tbig <- rbind(boot.Tcells,boot.PSV)

months <- c(6,12,18,24)

surfun <- function(csvname,endpoint,LOS,IDname,Nboot=2500,CIwidth=0.95){
  dat <- read.csv(csvname,na='.')
  dat <- dat[!is.na(dat$DV),]
  uID <- unique(dat[[IDname]])
  #print(uID)
  TTN <- c(); CEN <- c();
  for (j in 1:length(uID)){
    Tlit <- dat[is.element(dat[[IDname]],uID[j]),]
   # print(Tlit)
    if (nrow(Tlit)==0){print(paste(j,uID[j]))}
    igood <- which(Tlit$DV <= LOS)
    if (length(igood)>0){
      TTN <- c(TTN,Tlit$Days.PSO[min(igood)])
      CEN <- c(CEN,0)}
    else{
      TTN <- c(TTN,max(Tlit$Days.PSO))
      CEN <- c(CEN,1)
    }
  }
  Tout <- data.frame(uID,TTN,CEN)
  Tout$endpoint <- endpoint
  print(min(Tout$TTN))
  tsim <- seq(0,max(Tout$TTN),by=1)
  bootsfun = function(Npat){
    booti <- sample.int(Npat,size=Npat,replace=TRUE)  # which replicate to take for each patient
    Ttest <- Tout[booti,]
    ss <- survfit(Surv(TTN,1-CEN)~1,data=Ttest)
    ssim <- rep(NA,length(tsim))
    ssim[ss$time+1] = ss$surv
    ssim <- nafill(ssim,'locf')
    return(ssim)
  }
  
  Npat <- length(uID)
  foop <- replicate(Nboot,bootsfun(Npat))
  
  QQ <- rowQuantiles(1-foop,probs=c((1-CIwidth)/2,0.5,1-(1-CIwidth)/2))
  lower <- QQ[,1]
  median <- QQ[,2]
  upper <- QQ[,3]
  
  
  # sfit <- survfit(Surv(TTN,1-CEN)~1,data=Tout)
  # tsim <- sfit$time/dinm
  # median <- 1-sfit$surv
  # lower <- 1-(sfit$surv - 2*sfit$std.err)
  # upper <- 1-(sfit$surv + 2*sfit$std.err)
  tsim <- tsim/dinm
  Tsurv <- data.frame(tsim,lower,median,upper)
  Tsurv$endpoint <- paste(endpoint,'>',LOS)
  Tsurv$type <- 'observed'
  
  return(Tsurv)
}

sur.PSV <- surfun('../monolix/PSV-neut-titer.csv','PSV Neutralizing Titer',LOS=20,IDname='Donor.ID')
sur.Tcells <- surfun('../monolix/CD4-spike-T.csv','Spike-specific CD4+ Tcells (%)',LOS=0.03,IDname='UID')
sur.both <- rbind(sur.PSV,sur.Tcells)

bigdat <- rbind(sur.both,Tbig)

bigdat <- bigdat[bigdat$endpoint=='PSV Neutralizing Titer > 20',]

plo<-ggplot(data=bigdat) + geom_line(aes(x=tsim,y=median,color=type),size=1) + 
  geom_ribbon(aes(x=tsim,ymin=lower,ymax=upper,fill=type),alpha=0.25) +
  coord_cartesian(xlim=c(0,12))+ scale_x_continuous(breaks=seq(0,18,by=2)) +
  scale_color_discrete() +
  scale_y_reverse(labels=function(x){100-100*x}) +
  xlab('Months post symptom onset') + 
  ylab(paste0('% of population above LOS')) + 
  theme_bw() + theme(legend.position=c(0.9,0.9), legend.title = element_blank()) + #stat_ecdf(data=dat.both,aes(x=TTel,color=endpoint),size=2)
  #geom_label(data=Tbig[is.element(Tbig$tsim,months),],aes(x=tsim,y=median,color=endpoint,label=paste0(100-100*signif(median,2),'%'))) +
  facet_grid(rows=vars(endpoint)) 

print(plo)

# ggplot(data=Tbig) +  +
#   
#   geom_line(data=sur.both,aes(x=tsim/dinm,y=1-median,color=endpoint),size=1) + 
#   geom_ribbon(data=sur.both,aes(x=tsim/dinm,ymax=1-lower,ymin=1-upper,fill=endpoint),alpha=0.25) +
#   coord_cartesian(xlim=c(0,30))+ scale_x_continuous(breaks=c(6,12,18,24)) +
#   scale_color_discrete() +
#   scale_y_reverse(labels=function(x){100-100*x}) +
#   xlab('Months post symptom onset') + 
#   ylab(paste0('% of population above LOS')) + 
#   theme_bw() + theme(legend.position='none') + #stat_ecdf(data=dat.both,aes(x=TTel,color=endpoint),size=2)
#   geom_label(data=Tbig[is.element(Tbig$tsim,months),],aes(x=tsim,y=median,color=endpoint,label=paste0(100-100*signif(median,2),'%'))) +
#   facet_grid(rows=vars(endpoint)) 
```
