## Supplementary material for "Using mixed-effects modeling to estimate decay kinetics of response to SARS-CoV-2 infection": R codes, analysis data set, and monolix files: README.txt

supplementary materials for Bottino et al BIOXRIV submission:
Using Mixed Effects Modeling to estimate decay kinetics of response to SARS-CoV-2 infection
No warranty
Attribution CC-BY-ND as inherited by BIORXIV publication
Keep the directory structure if you want the codes to work properly. 
Run the monolix first, then the R code
The monolix dataset is derived from Dan et al (Crotty) Science, Immunological memory to SARS-CoV-2 assessed for up to 8 months after infection, 2021.
